# Beav: A bacterial genome and mobile element annotation pipeline

**DOI:** 10.1101/2024.01.25.577299

**Authors:** Jewell M. Jung, Arafat Rahman, Andrea M. Schiffer, Alexandra J. Weisberg

## Abstract

Comprehensive and accurate genome annotation is crucial for inferring the predicted functions of an organism. Numerous tools exist to annotate genes, gene clusters, mobile genetic elements, and other diverse features. However, these tools and pipelines can be difficult to install and run, be specialized for a particular element or feature, or lack annotations for larger elements that provide important genomic context. Integrating results across analyses is also important for understanding gene function. To address these challenges, we present the Beav annotation pipeline. Beav is a command-line tool that automates the annotation of bacterial genome sequences, mobile genetic elements, molecular systems and gene clusters, key regulatory features, and other elements. Beav uses existing tools in addition to custom models, scripts, and databases to annotate diverse elements, systems, and sequence features. Custom databases for plant-associated microbes are incorporated to improve annotation of key virulence and symbiosis genes in agriculturally important pathogens and mutualists. Beav includes an optional *Agrobacterium*-specific pipeline that identifies and classifies oncogenic plasmids and annotates plasmid-specific features. Following the completion of all analyses, annotations are consolidated to produce a single comprehensive output. Finally, Beav generates publication-quality genome and plasmid maps. Beav is on Bioconda and is available for download at https://github.com/weisberglab/beav.

## Introduction

Correct and comprehensive genome annotation is critical for characterizing and understanding microbial function and evolution. However, *de novo* annotation of under-studied microorganisms can be challenging. Gene names and proteins may be poorly annotated, and representative sequences of these taxa are often not integrated into databases of existing annotation tools^1^. Bacteria with clinical relevance are overrepresented in gene and genome databases, and important virulence genes from these organisms can be confidently identified and annotated^2^. However, annotation of many phytopathogens and plant-associated microbes often fails to identify and name key genes important for plant-microbe interactions, symbiosis, and virulence. For instance, effector proteins secreted by phytopathogens have a fundamental role in the plant disease process, yet they are underrepresented in annotation tools and databases^3^. While these key virulence loci may be characterized and classified in species-specific databases and publications, these annotations are often not incorporated into annotation databases or current tools.

Additionally, current whole-genome annotation pipelines typically focus on the annotation and function of individual genes, but often do not report information about their genomic context. Genes may be part of biosynthetic gene clusters or metabolic pathways, be organized into operons, represent loci encoding larger macromolecular structures, or be carried on mobile genetic elements (MGEs) and/or prophage elements. Gene regulatory elements, such as promoters or transcription factor binding sites, are important for understanding how genes are expressed, but these elements are also typically not annotated. Understanding the genomic context and regulation of genes is crucial for understanding their function.

Mobile genetic elements, such as plasmids, integrative and conjugative/mobilizable elements (ICEs/IMEs), integrons, prophages, and other elements can be mobilized from cell to cell and play a major role in the horizontal transfer of genes between bacteria^4^. MGEs may be integrated into the chromosome, such as in the case of ICEs and transposons, or they may replicate independently, such as plasmids. MGEs can be incredibly diverse and vary in structure and function. This can make their identification and annotation difficult, especially in draft genome assemblies. The horizontal transfer of MGEs, either directly via conjugation, or indirectly on other elements, enable bacteria to rapidly respond to changes in their environment and acquire new traits that may be selectively advantageous^4–6^. For example, plasmids are associated with the movement and transfer of antimicrobial resistance (AMR) genes and genes associated with pathogenicity or mutualist symbioses^7–9^. ICEs and IMEs are also recognized as drivers of HGT and have been attributed to the spread of genes involved in AMR, pathogenesis, and symbiosis in diverse microbial taxa^10–12^. Integrons are genetic elements that can acquire and shuffle the order of gene “cassettes” relative to a single promoter^13^. The order of these genes can be rearranged in response to stress conditions, altering their expression^14^. Many integron gene cassettes have been found to encode for traits associated with virulence, resistance, and host-microbe interactions^15^.

Bacterial genomes often encode for one or more secretion systems. These secretion systems are diverse multi-protein complexes that have a range of functions, including those that play a fundamental role in the conjugative transfer of MGEs, interbacterial communication and competition, and/or host-microbe interactions^16–18^. Conjugation of plasmids and ICEs is typically facilitated by a type IV secretion system (T4SS)^19, 20^. Type IV secretion systems sometimes provide other functions beyond conjugation. One of the most well-studied T4SS is in the phytopathogen *Agrobacterium tumefaciens*, where it facilitates the inter-Kingdom transfer of DNA^21^. Microbial defense systems are also encoded on and transferred via MGEs^22^. These diverse systems provide defense against invading DNA, either from phage or plasmids. With an abundance of MGEs in bacteria and genome data, characterizing MGEs and the systems they interact with is essential to answer questions regarding microbial evolution, ecology, and resistance. Despite the importance of MGEs, few tools exist to comprehensively characterize MGEs within bacterial genomes^23^.

Comprehensive characterization of specific genetic regions of interest requires multiple genome annotation tools. Many independent tools for the annotation of specific genetic systems have recently been published^24–28^. However, the results of these separate analyses must be combined to get a full picture of gene function and context. Additionally, the installation and use of these tools presents a challenge, even for those with computational proficiency. Installation of individual annotation software tools can be laborious and complicated by numerous or conflicting dependencies. This, coupled with unclear or minimal documentation, can limit the accessibility of powerful tools. Moreover, genome analysis with multiple annotation tools also requires manual parsing and cross-correlating numerous output files, which requires experience with the command line.

In response, we present Beav, a command-line tool that streamlines and automates bacterial genome and mobile genetic element annotation. The Beav pipeline incorporates multiple annotation tools, automating the process of running, parsing, and combining results into a single easy-to-read output. The Beav pipeline also includes several tools and databases that enhance the annotation of plant-associated microbes, including genes and regulatory elements important to phytopathogens and mutualist symbionts. Additionally, an optional *Agrobacterium*-specific pipeline identifies the presence of oncogenic Ti and Ri plasmids and classifies them under a published scheme^29, 30^. This pipeline also annotates Ti/Ri plasmid-specific regulatory elements and reports the taxonomic classification of the input strain under the *Agrobacterium* biovar/genomospecies scheme^31^. Finally, Beav generates a visualization of the position of annotated gene clusters and mobile elements in the genome. Additionally, Beav can generate a separate plot to visualize oncogenic Ti/Ri plasmids, if present.

Beav is a comprehensive genome annotation pipeline for bacteria and associated mobile genetic elements. Beav and its dependencies are available for installation via conda and requires minimal user input for installation and usage. The pipeline uses pre-existing annotation tools in combination with custom scripts and databases to automate annotation and combines results in a single easy to read GenBank and/or GFF3 format output. Beav databases and source code are also freely available for download on GitHub at http://github.com/weisberglab/beav.

## Materials and Methods

### Usage of the Beav pipeline

Beav is designed to be user-friendly and includes multiple checks that verify the correct installation of dependencies and valid input arguments before running the pipeline. Beav is a command-line tool written in Python and shell script that runs on Unix-based operating systems such as Linux. The Beav pipeline will clearly indicate if prerequisites are installed correctly, skipped, or causing an error. Each annotation step of the pipeline is listed as it runs and labeled as ‘Done’ when complete. Tables and log files summarizing the output of each annotation program are also produced during the run. The Beav pipeline workflow includes multiple databases and annotation tools (**Figure 1**). Beav is packaged in Bioconda for ease of installation^32^. Installing Beav via conda will also install all required dependencies in a single environment. A separate program included with Beav will also automatically download and format all databases needed by Beav and its dependencies. Most steps in the workflow are optional and can be skipped. Users input a single file with the nucleotide sequence of their genome assembly in fasta format, along with any other optional parameters. Beav then annotates the genome and proceeds to run other annotation programs. Finally, results of each program are parsed, and the initial annotations for each gene are supplemented with information from each of the annotation tools and reported in GenBank and GFF3 format output files. Regions representing mobile genetic elements and gene regulatory elements are also annotated in the final output files. If a Beav run is interrupted or the user wishes to re-run the analysis with additional tools, Beav can also be restarted with the “--continue” option to finish any incomplete analyses.

**Figure 1.**
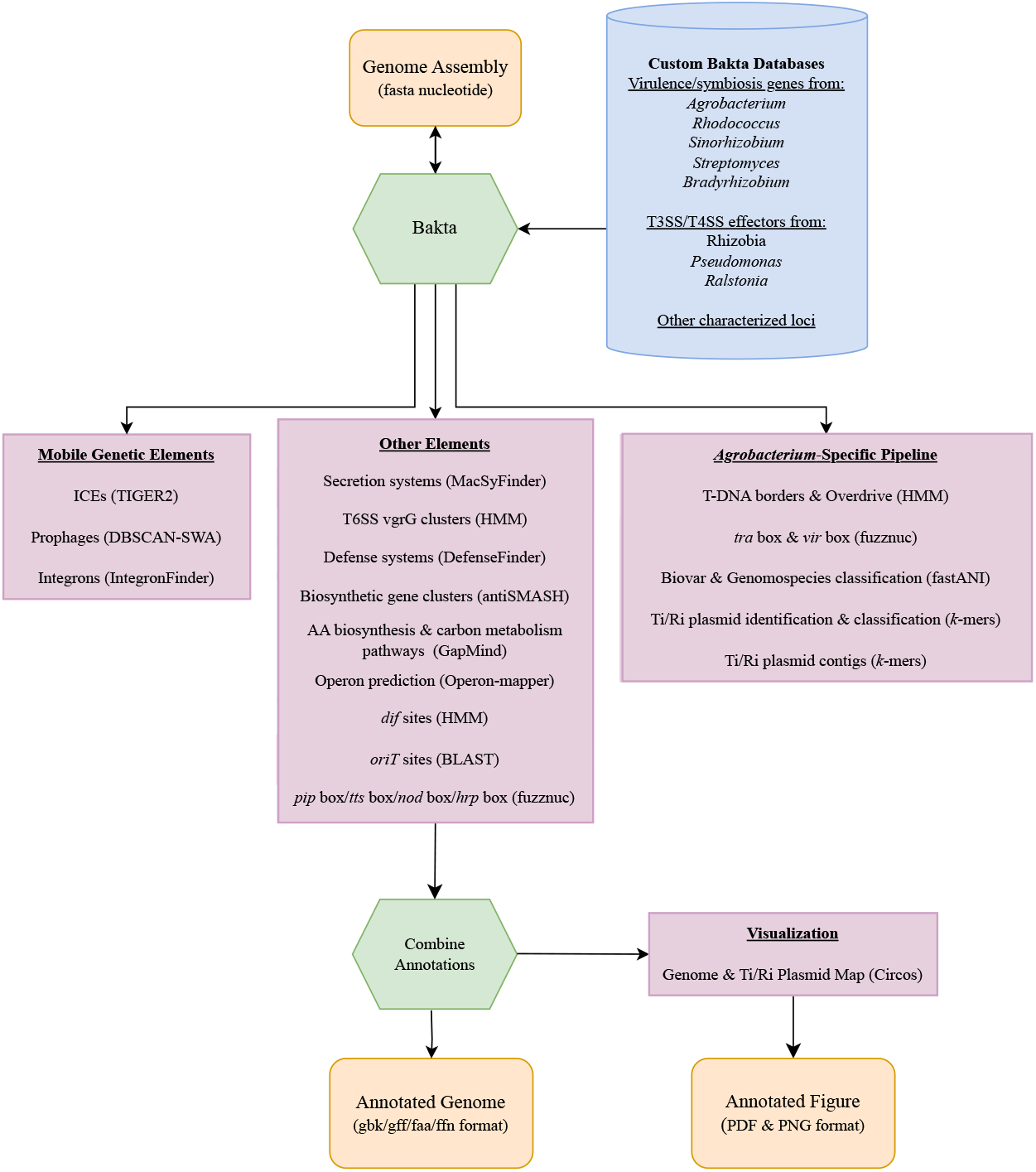
Summary of the Beav pipeline workflow. Beav takes a fasta nucleotide file as input and uses Bakta with a custom gene database to generate preliminary annotations. Following this, the pipeline runs several suites of annotation programs to annotate mobile genetic elements and other systems and gene clusters in the genome. An optional *Agrobacterium*-specific pipeline can be run that identifies and classifies Ti/Ri plasmids and annotates features specific to these elements. Finally, annotations are combined and output into standard bioinformatic file formats. Visualizations of annotations across the whole genome as well as Ti/Ri plasmids are generated in raster and vector image formats.

### Initial annotation and custom gene databases

Beav takes as input an assembled genome in fasta nucleotide format. Bakta is used for the initial annotation of genes and other loci^33^. The Bakta run is supplemented with custom Bakta-formatted gene databases that enrich the annotation of plant-associated microbes. Beav incorporates several custom databases into Bakta to ensure the correct names of virulence and symbiosis genes for numerous phytopathogens and mutualists (**Table 1**). These databases were compiled from published datasets and online resources^10, 29, 30, 34–38^. Databases include known virulence genes and genes encoding secreted effectors from phytopathogenic *Agrobacterium tumefaciens*, *Pseudomonas syringae*, *Ralstonia solanacearum*, *Rhodococcus fascians*, *Streptomyces scabiei*, and key loci from mutualist *Bradyrhizobium*, *Rhizobium*, and *Sinorhizobium*.

**Table 1:** Custom Bakta annotation databases for plant-associated microbes.

| Organism | Genes and loci | Reference |
| --- | --- | --- |
| *Agrobacterium* | *vir*, *GALLS*, *acc*, agrocin84 biosynthesis, opine synthases, T-DNA oncogenes, *trb/tra*, *upp* attachment cluster | 29,30,85 |
| *Bradyrhizobium* | *nod*, *nol/nop*, *nif/fix* symbiosis genes, Type III secreted effectors | 10 |
| *Rhodococcus* | *fas/att* virulence genes | 34,35 |
| *Sinorhizobium* | *nod*, *nol/nop*, *nif/fix* symbiosis genes | *Sinorhizobium fredii* HH103 (NCBI: GCA_000283895.1) |
| *Streptomyces* | *txt*, *nec1*, *tomA*, *fas* virulence genes | 36 |
| *Ralstonia* | Type III secreted effectors | 37 |
| *Pseudomonas* | Type III secreted effectors | 38 |

Following the Bakta run, the Beav pipeline annotates promoters and binding sites associated with plant-associated microbes, including *pip* box (plant-induced promoter), *tts* box and *hrp* box (regulating genes associated with type III secretion systems), *nod* box (regulating genes associated with nodulation in nitrogen fixation symbiosis), *tra* box (regulation of conjugation loci of Ti/Ri plasmids), and *vir* box (regulation of Agrobacterium virulence genes) elements. For several of these elements, the EMBOSS program fuzznuc is used to identify elements based on conserved sequence patterns^39^. The *pip* box promoter is identified with the pattern “TTCGBN(15)TTCGB”^40^. The *hrp* box promoter is identified with the consensus pattern “GGAAC[CT]N(15,17)CCACNNA”^41–43^. The HMMER3 suite program nhmmer is used to search with custom HMM profiles that we have built for other elements from known sequences, including *nod* box and *tts* box regulatory elements, and chromosome *dif* sites^10, 44–51^. Bedtools is used to ensure that predicted regulatory sequences do not overlap with coding sequences^52^.

### Annotation of mobile genetic elements

The Beav pipeline includes several tools to annotate diverse mobile genetic elements in genomes, including ICEs, integrons, and prophage elements. TIGER2 is used to identify and annotate monopartite ICEs integrated into the genome^28^. The boundary regions of the identified ICEs, as well as the integrase and predicted target genes and sequences (*attB*, *attP*), are reported in a “mobile_element” feature in the final annotation. Plasmid/ICE origin of transfer (*oriT*) sites are annotated using a database of known *oriT* sequences subset to remove duplicate and very short sequences^53^. Blastn with the options “-task blastn-short -outfmt ‘6 std qlen slen qseq sseq’ -dust no -qcov_hsp_perc 20” is used to identify putative *oriT* sequences in the input assembly^54^. Blast hits are further filtered to those with an e-value of 0.1 or less and a minimum alignment length of 20 bp. DBSCAN-SWA is used to annotate prophages integrated into the genome^27^. A mobile_element feature reporting the entire prophage region and classification is also included in the final annotation. IntegronFinder is used to identify and annotate integron loci and cassettes^26^. Predicted integron regions are annotated, including the integrase *intI* gene, promoter, and the location of *attC* sequences bordering cassettes.

### Annotation of other systems and functional loci

Beav incorporates additional annotation tools that identify other conserved gene clusters or provide further context for gene function. MacSyFinder with the TXSScan models is used to identify genes and gene clusters encoding for diverse secretion systems^24, 55^. DefenseFinder is used to characterize the presence of microbial defense systems^56^. These systems provide defense against invasion by foreign DNA, including phage and plasmids^57^. AntiSMASH is used to identify and annotate biosynthetic gene clusters^25^. GapMind is used to associate genes with amino acid biosynthesis pathways as well as genes involved in the catabolism of small carbon metabolites^58, 59^. Genes in bacterial genomes are often encoded in and expressed as single transcriptional units called operons^60^. Beav can optionally submit jobs to the operon-mapper webserver, download completed results, and parse the resulting output^61^. Operon tags associating genes to specific predicted operons are added to gene features in the final annotation. If the operon pipeline is run, Beav will also annotate type VI secretion system *vgrG* clusters. These gene clusters encode for the vgrG spike protein as well as adapters, toxins, and cognate immunity genes^62^. The HMMER3 program nhmmer is used to search with HMM profiles for vgrG (TIGRFAMs TIGR01646.1 and TIGR03361.1). Operons containing *vgrG* genes are then reported as putative *vgrG* clusters.

### *Agrobacterium*-specific pipeline

In addition to generic annotation tools applicable to all bacteria, Beav includes an optional pipeline for annotating features specific to the phytopathogen and genetic engineering tool *Agrobacterium tumefaciens*. Using the optional *Agrobacterium* pipeline, *Agrobacterium* genomes can be further annotated with Ti/Ri plasmid-specific loci, including T-DNA borders, overdrive, *vir* box, and *tra* box elements. Fuzznuc with the pattern “RTTDCAWWTGHAAY” is used to annotate the *vir* box virulence gene promoter^63^. The Ti/Ri plasmid conjugation loci promoter *tra* box is identified by the consensus pattern “WNGTGMARAWYTGCACDW”^64–66^. The HMMER3 suite program nhmmer is used to search with custom HMMs that we developed based on the sequence of known T-DNA border and overdrive sequences^29, 51, 67–69^. Beav also automates the identification of oncogenic Ti and Ri plasmid sequences in the input genome assembly and classifies the plasmid type based on a classification scheme^29, 30^. The BBTools program comparesketch.sh is used to identify Ti/Ri plasmid contigs based on a custom database of known oncogenic plasmids^70^. Output is reported as the presence of a Ti/Ri plasmid, its classification, and a list of contigs associated with that plasmid. FastANI and a database of representative *Agrobacterium* taxa is used to classify the input *Agrobacterium* strain into biovar and genomospecies-level designations^31, 71^. Beav also includes extra scripts to run the *Agrobacterium* analyses independent of the full annotation pipeline.

### Parsing and combining annotations

While running multiple annotation tools can be informative, cross correlating the results of multiple analyses and annotation tools can be challenging. Results are often present in multiple files, and in different formats. Beav solves this issue by automatically parsing the results of each step of the pipeline and incorporating those results into a single output file. A custom Python script and the BioPython library is used to parse the Bakta GenBank output and the results of each step of the pipeline, and add information as new loci or to associated gene features^72^. In the GenBank output, mobile genetic elements and prophage are added as “mobile_element” features, and regulatory and other elements are labeled as “misc_feature” elements. Additional information about genes and coding sequences is added as “notes” in the feature qualifiers field of existing features. The annotation software used to make each inference is also listed. Final annotations are output in GenBank (gbk) and GFF3 formats, and coding sequences are output in fasta nucleotide (ffn) and amino acid (faa) formats. Output files of each tool are organized into directories for easy navigation of results and annotations. This alleviates tedious navigation of multiple output files and keeps results organized. Logs and table format output of each tool are also produced and stored in relevant folders. At the end of a Beav run, all software tools used in the analysis, along with their versions and suggested citations, are listed for ease of reporting results in publications.

### Visualization of genome and plasmid annotations

Data visualization is important for understanding large scale genome structure and organization. However, converting data into the correct format for existing tools and generating publication-quality visualizations can be difficult. Beav automates this process and generates figures of major annotated features and gene clusters across the genome (**Figure 2**). The Python package PyCirclize is used to generate Circos plot visualizations^73^. Beav generates Circos plots that visualize whole genome structure, including contigs, the position of secretion system-encoding loci along with their types (i.e. T3SS, T4SS, T6SS, etc.), ICEs, prophage elements, specialized metabolite gene clusters, and tRNA/rRNA position. If Beav is run with the optional *Agrobacterium*-specific pipeline and a Ti or Ri plasmid is detected, a second Circos plot is generated that visualizes Ti/Ri plasmid contigs and associated features (**Figure 3**). This plot shows the structure and content of the Ti/Ri plasmid and key loci, including virulence genes, T-DNA regions and borders, plasmid replication (*repABC*) and conjugation (*tra*/*trb*) loci, specialized nutrient (opine) synthase and transport genes, and regulatory elements such as the *tra* box and *vir* box. Beav also includes a supplemental stand-alone command-line tool to visualize other GenBank files.

**Figure 2.**
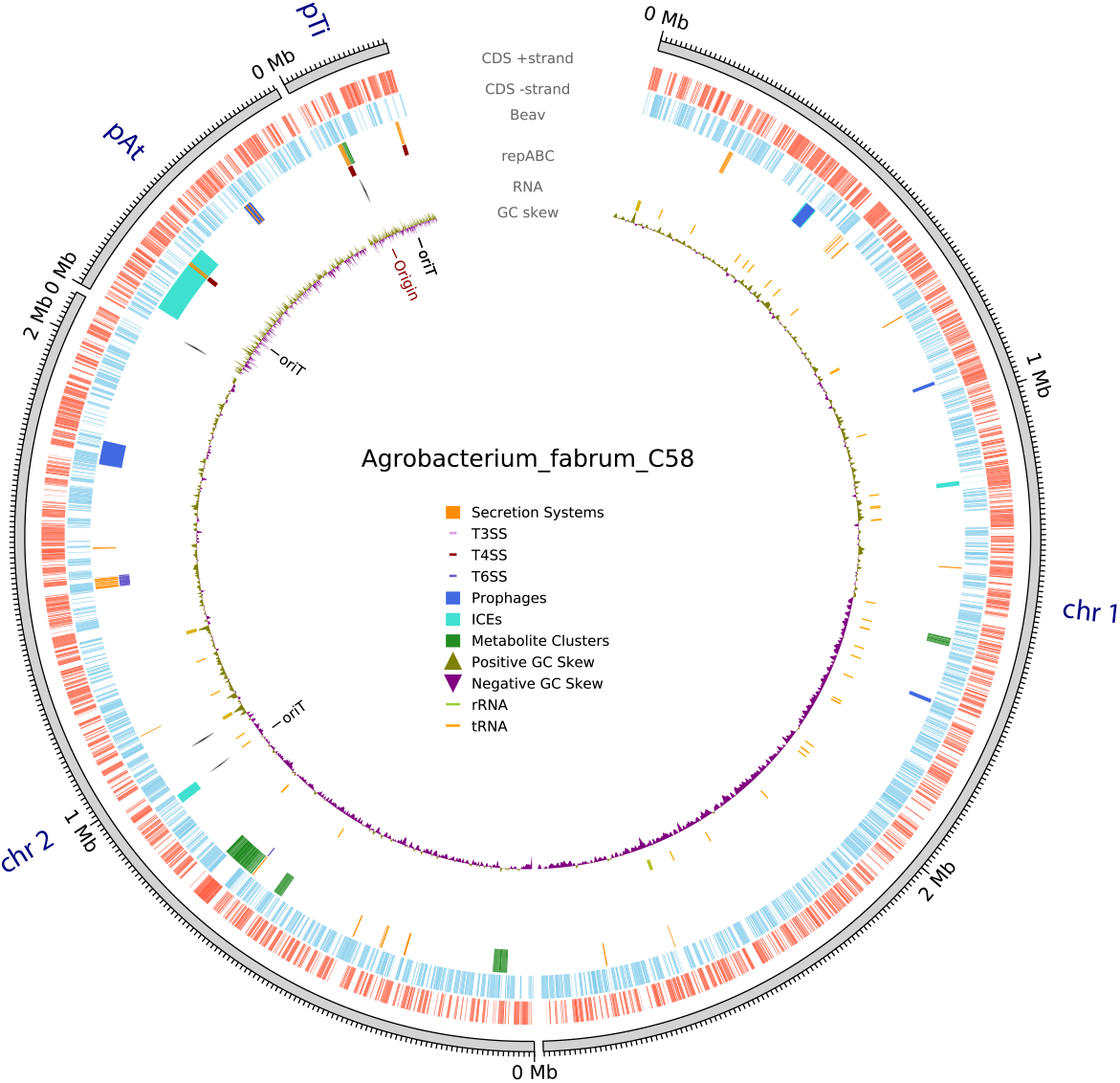
Example Circos plot of whole genome annotations automatically generated by Beav. There are 6 inner tracks showing the position of different elements and features. The outer-most track shows the length of each assembly contig. The next two tracks indicate CDS position on the +/- strand. The third track shows elements annotated by the Beav pipeline, including secretion systems, secondary metabolite gene clusters, and mobile genetic elements. The specific type of secretion system, such as T3SS, T4SS, or T6SS, is plotted as a small colored rectangle along the main feature. Regions encompassing MGEs, such as ICEs and prophages, are indicated as colored blocks. The next track indicates the position of plasmid and secondary chromosome replication loci. The following track indicates the position of ribosomal RNA and other RNA elements. The next track indicates GC Skew across a contig. The innermost track includes markings indicating the position of origin of replication (*oriC*) and MGE origin of transfer (*oriT*) sites.

**Figure 3.**
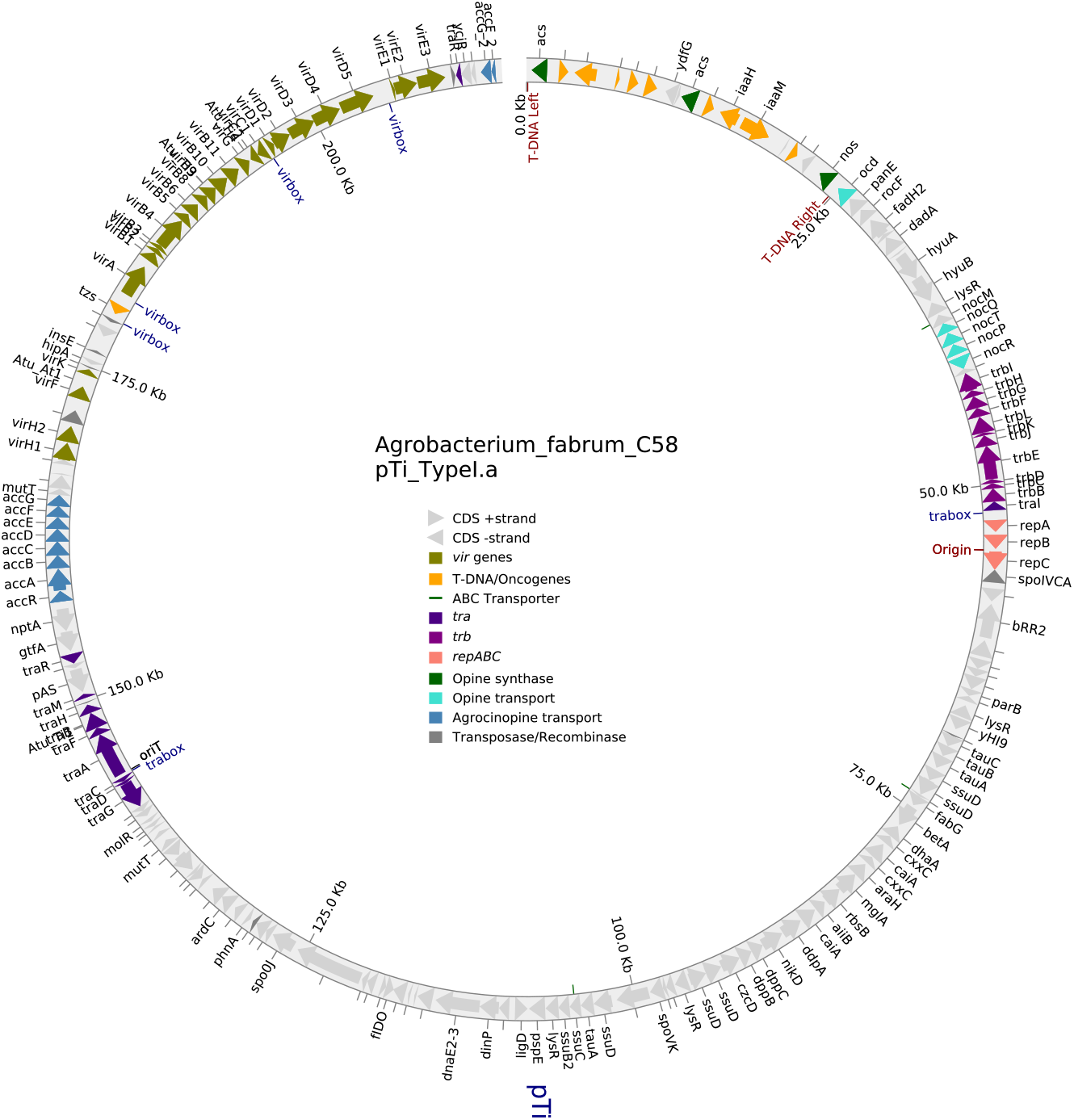
Example Circos plot visualizing oncogenic Ti/Ri plasmids as generated by the optional *Agrobacterium*-specific pipeline. Beav includes an optional *Agrobacterium*-specific pipeline. If this pipeline is run, an additional figure is generated showing contigs identified as representing Ti or Ri plasmids and their annotations. Genes and their strand are visualized as arrows. The presence of key plasmid loci, including T-DNA genes, virulence (*vir*) genes, specialized nutrient (opine) synthesis and catabolism loci, and plasmid replication (*repABC*) and conjugation (*trb*/*tra*) are indicated by arrow color. Other plasmid and regulatory elements, such as T-DNA borders, *vir* box, *tra* box, and the position of ABC transporter genes are indicated. Transposases, integrases, and IS element-associated loci are colored dark gray. All other genes are colored light gray. Gene names, where present, are visualized outside of the ring. If run on draft assemblies with multiple Ti/Ri plasmid contigs, all will be included in the visualization. Several text labels have been adjusted for visibility, otherwise this figure appears as generated by Beav.

## Results

### Comparison of annotations across pipelines

To test the performance and annotation improvements of Beav relative to Bakta alone and the RefSeq PGAP pipelines, we annotated a diverse set of representative bacterial genomes, including *Agrobacterium fabrum* C58 (NCBI: GCA_000092025.1), *Escherichia coli* 131 (NCBI: GCA _005221985.1), *Pseudomonas syringae* DC3000 (NCBI: GCF_000007805.1), *Rhodococcus fascians* D188 (NCBI: GCF_001620305.1), and *Aeromonas caviae* WP2-W18-ESBL-01 (NCBI: GCF_014168635.1). We then compared the number of genes annotated as “hypothetical protein” and “uncharacterized protein” by the three pipelines (**Table 2**). For the Beav pipeline, annotation of the *Agrobacterium fabrum* C58 genome was run with the optional *Agrobacterium* pipeline while all other genomes were run with default options. For this test, a gene is considered annotated by Beav if a “note” was added to loci annotated as a hypothetical or uncharacterized protein. Overall, Beav and Bakta predicted a function for more genes than RefSeq PGAP. Beav and Bakta produced comparable numbers of annotated hypothetical proteins for all the genomes, which can be attributed to the Beav pipeline using Bakta for preliminary gene annotations. However, Beav consistently produced additional annotations for genes that Bakta annotated as hypothetical. These results demonstrate that the combined analyses of the Beav pipeline can improve genome annotation over a single tool.

**Table 2.**
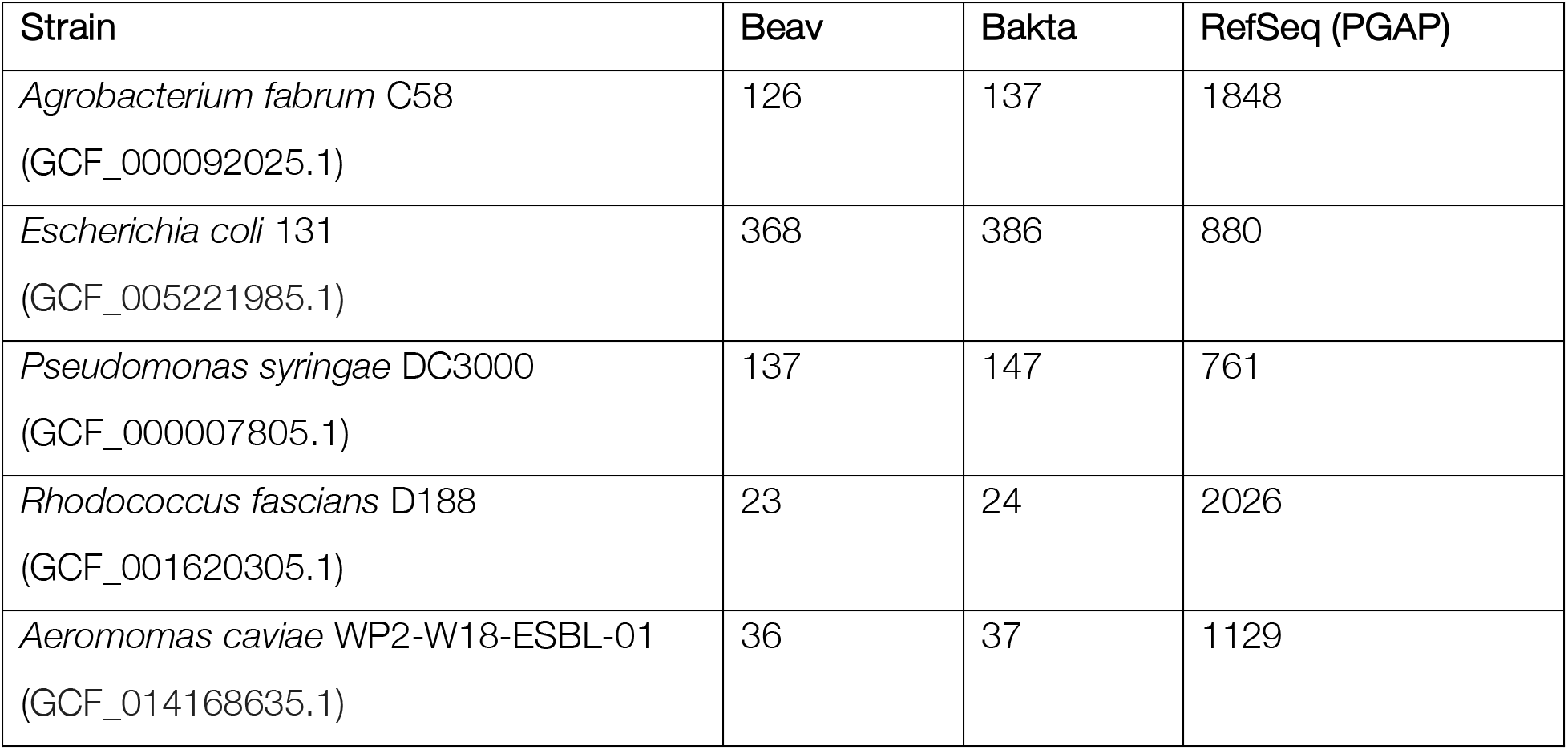
Counts of hypothetical and uncharacterized proteins in annotations produced by Beav, Bakta, and RefSeq PGAP.

| Strain | Beav | Bakta | RefSeq (PGAP) |
| --- | --- | --- | --- |
| *Agrobacterium fabrum* C58<br>(GCF_000092025.1) | 126 | 137 | 1848 |
| *Escherichia coli* 131<br>(GCF_005221985.1) | 368 | 386 | 880 |
| *Pseudomonas syringae* DC3000<br>(GCF_000007805.1) | 137 | 147 | 761 |
| *Rhodococcus fascians* D188<br>(GCF_001620305.1) | 23 | 24 | 2026 |
| *Aeromonas caviae* WP2-W18-ESBL-01<br>(GCF_014168635.1) | 36 | 37 | 1129 |

To further assess the usefulness of annotations produced by the Beav pipeline, we summarized the total number of systems, mobile elements, gene clusters, and new features in annotations of diverse bacteria (**Table 3**). For each test genome, Beav successfully identified the genes of multiple complete biosynthetic gene clusters and defense systems. Further, at least one putative secretion system was identified in each genome. MGE annotations varied based on the analyzed strain. For instance, several ICEs were found in the *A. caviae*, *A. fabrum*, *E. coli*, and *P. syringae* genomes, while zero were found in *R. fascians*. However, these elements might not be present or common in certain bacterial lineages or strains. These results indicate that the Beav pipeline can annotate complete systems and genetic elements applicable to a broad range of bacterial taxa.

**Table 3.**
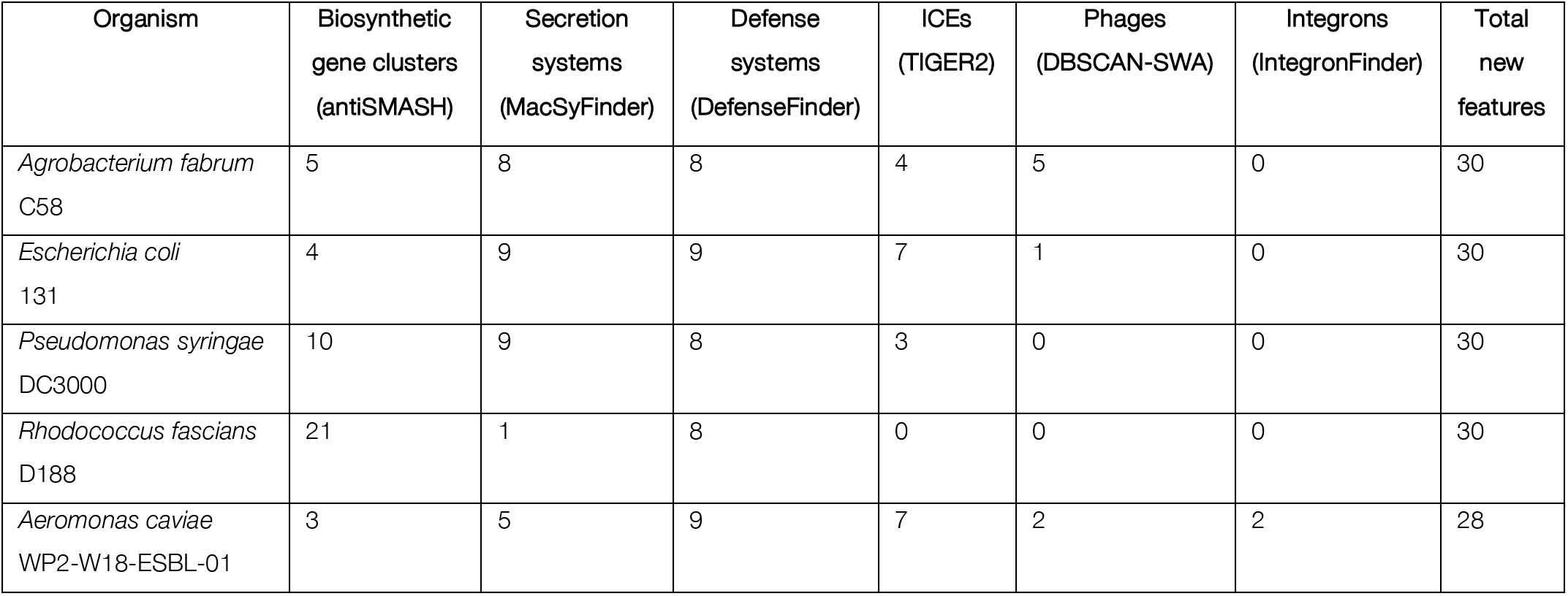
Summary of annotated features and systems predicted by Beav for diverse bacteria.

### Analysis of Beav pipeline runtime

To evaluate the runtime of the Beav pipeline, we summarized the amount of time it took Beav and the component steps to run to completion for a diverse set of genomes. Three replicates were run on a cluster computer using eight cores of an AMD EPYC 7601 processor running at 2.2 GHz per core, with 512 GB of available RAM. The average run time of the overall pipeline and each subcomponent was calculated (**Table 4**). In almost all cases, Beav took less than one hour to run the entire pipeline. The longest steps included the TIGER2 ICE analysis and Bakta which both took 18 minutes on average to run. TIGER2 runtime increased with the number of ICEs encoded in the genome. For example, *R. fascians* D188 encodes 0 ICEs and TIGER2 took about three minutes to run, while *A. fabrum* C58 encodes 4 ICEs and TIGER2 took 36 minutes to run. This indicates that the TIGER2 analysis will not greatly increase the run-time if no ICEs are present. However, for genomes that contained multiple ICEs, TIGER2 analysis is a major proportion of Beav runtime. Other steps completed very quickly and did not add much to overall runtime. The other program’s run times vary based on the features in the genome, but their analysis did not individually exceed five minutes. For example, DefenseFinder, MacSyFinder, and GapMind took around one and a half minutes or less to run for each test genome. These results show that while the Beav pipeline does take longer to run than Bakta, it is not unreasonably longer for the additional annotations Beav provides. Individual programs or steps in the Beav pipeline can be skipped to reduce computational needs or shorten runtime if necessary.

**Table 4.** Runtime for Beav annotation pipeline components.

| Organism | Beav total<br>run time<br>[hh:mm:ss] | Bakta | MacSyFinder | DefenseFinder | antiSMASH | DBSCAN-SWA | GapMind | TIGER2 | IntegronFinder |
| --- | --- | --- | --- | --- | --- | --- | --- | --- | --- |
| *Agrobacterium fabrum*<br>C58<br>(GCF_000092025.1) | 1:04:29 | 00:16:20 | 00:00:54 | 00:01:16 | 00:02:05 | 00:02:07 | 00:01:25 | 00:36:53 | 00:00:32 |
| *Escherichia coli*<br>131<br>(GCF_005221985.1) | 00:53:50 | 00:18:25 | 00:00:50 | 00:00:41 | 00:01:42 | 00:01:28 | 00:01:11 | 00:26:23 | 00:00:31 |
| *Pseudomonas syringae*<br>DC3000<br>(GCF_000007805.1) | 00:53:24 | 00:19:26 | 00:00:57 | 00:00:42 | 00:03:06 | 00:01:02 | 00:01:14 | 00:23:35 | 00:00:48 |
| *Rhodococcus fascians*<br>D188<br>(GCF_001620305.1) | 00:34:24 | 00:18:18 | 00:1:03 | 00:01:55 | 00:04:22 | 00:00:36 | 00:01:33 | 00:03:25 | 00:01:54 |
| *Aeromonas caviae*<br>WP2-W18-ESBL-01<br>(GCF_014168635.1) | 00:51:34 | 00:17:20 | 00:01:10 | 00:01:08 | 00:01:21 | 00:01:11 | 00:01:06 | 00:22:36 | 00:03:43 |

## Discussion

Comprehensive genome annotation can be challenging since it requires the use of multiple tools and analyses, some of which are difficult to install or run. It also requires parsing and interpreting the results of these tools, and cross-correlating results for individual genes and loci. Tools can be broadly separated into general genome annotation pipelines, which identify open reading frames and annotate gene and RNA function, and more specialized programs that identify and characterize specific kinds of loci or elements, such as secretion systems or ICEs. Most published annotation pipelines focus on gene identification and annotation and leave the characterization of other kinds of loci to other tools. Few tools combine both kinds of analysis into a single pipeline. Beav is designed to fill this gap and merge results from both general and specialized annotation tools, while adding additional annotations available in no other program.

Several current genome annotation pipeline options include web-based tools, such as BacAnt, DFAST, Galaxy/Apollo, NCBI PGAP, and RAST^74–78^. Web-based services such as Galaxy and Apollo address many of the challenges of annotation by providing a web-based platform that integrates multiple popular annotation tools into a dashboard, making complex analysis and generating figures easier^75, 76^. These web services make genome annotation available to users with little to no command-line experience. However, the analysis of whole genome sequences can be computationally intensive, precluding web servers from high throughput analysis or providing more comprehensive analyses beyond gene identification. Users are often limited to a small number of concurrent jobs, and job wait times can be extensive. This can restrict webserver use to analyses of a relatively small number of genomes.

Command-line pipelines are targeted for high-throughput projects and users with some computational experience. There are several established and published command-line annotation pipelines available, including Bakta, Prokka, and MicrobeAnnotator^33, 79, 80^. These tools tend to specialize for identification of open reading frames encoding for genes or RNA loci and annotate their predicted function using databases of known genes. Tools such as Bakta work very well for this process and provide high quality annotations for individual genes. Rather than reimplementing this step, we relied on Bakta for initial gene annotations in Beav. We then complement the Bakta annotations with other tools and scripts. However, complex analyses that involve more than one software tool quickly become complicated. Each program produces several outputs and results, and it can be difficult to navigate through every output file. Manually parsing these outputs for valuable information is a time-consuming process. We designed Beav to alleviate the challenges of complex genome annotation and provide a comprehensive tool that users with a basic level of command-line experience can use. The Beav pipeline makes existing annotation software more accessible. Installation of the Beav pipeline and all its dependencies is made easy using Conda. Conda manages installation so that each program does not need to be installed individually and users do not have to manage dependencies. Likewise, Beav includes a tool that downloads and installs databases and updates for each of the prerequisite programs. Beav was developed with a highly customizable workflow that allows for programs to be run sequentially, independently, or skipped depending on the user’s needs. This makes using existing programs easier since running the program is automated and does not require intensive knowledge of each program’s usage. Additionally, the pipeline parses the results of multiple annotation tools and combines them into one output. This makes interpretation of annotations easier as results from several programs are merged.

Having access to these powerful tools makes Beav applicable across many fields of bacterial research. Beav also includes specific databases and tools for improving the quality of annotations for plant-associated microbes, particularly agriculturally relevant phytopathogens and symbiotic mutualists. Consistent annotation of virulence genes and their names, such as those encoding for secreted effector proteins, is important for effective communication and understanding of pathogens. This makes Beav a valuable tool for plant-microbe and phytopathogen-related studies as the pipeline has gene databases and novel tools that provide reliable naming and annotation conventions. There is currently no single database listing names and representative sequences for virulence genes or effectors for all plant pathogens, and research communities of different pathogens have different standards and naming conventions. Future updates to the Beav database could include genes from other plant-associated microbes, such as additional phytopathogens and mutualists. Unlike other annotation tools, Beav detects promoter and regulatory features unique to plant-associated microbiota. Methods for annotating known promoter and regulatory regions exist and require manual input of patterns or custom models for each region. With Beav, annotation of these elements is automated and greatly reduces the workload that would come with characterizing these regions individually. Accurate and comprehensive whole genome annotation of phytopathogens and maintaining a reliable repository of genes important for plant-microbe interactions is crucial for pathogen management. We developed Beav to address the need for bioinformatics tools that assist in the analysis of plant-associated microbes and minimize the naming errors commonly associated with these taxa. However, the Beav pipeline and its incorporated tools are broadly applicable to diverse taxa of bacteria.

Beav is the only tool that features an *Agrobacterium*-specific pipeline developed to offer a standardized method for *Agrobacterium* genome annotation. *Agrobacterium* is both an economically important plant pathogen and a biotechnology tool for genetically modifying plants. Thus, this sub-pipeline can aid in *Agrobacterium* pathology and genomics studies, as well as the development of new strains for use in plant transformation and engineering. The *Agrobacterium* oncogenic plasmids, the Ti and Ri plasmids, are diverse in both sequence and content^29, 30^. Identifying the presence of an oncogenic plasmid in a genome assembly, and correctly classifying its type, can be difficult, especially in draft assemblies. Consistent classification of plasmids can improve communication on differences between *Agrobacterium* strains. Beav fully automates this process and provides additional annotations for other key virulence elements and regulatory features. The custom gene database also provides for consistent naming of key oncogenic plasmid genes, some of which are frequently misannotated in other tools.

To extend the function of the current Beav pipeline, further work improving annotations and adding other kinds of genome features is still needed. Knowing operon structure is important for understanding gene expression and function, yet associating genes to operons is surprisingly difficult. In our experience, the most accurate *de novo* operon prediction tool that does not require RNA-Seq data is the operon-mapper webserver^61^. The current version of Beav can submit jobs remotely to this tool. However, using webservers requires an internet connection, limits concurrent jobs, affects version controlling, and webservers may go down or be no longer supported. We hope to adjust the current method of web-based operon mapping towards a local command-line tool, reducing wait times and the potential for network connection and server errors.

Bacterial secreted proteins play a major role in host-microbe interactions for phytopathogenic and clinically relevant bacteria^81^. Predicting and identifying genes encoding for secreted proteins is essential for understanding virulence. While Beav annotates genes with similarity to known secreted effectors, it currently does not identify novel effectors *de novo*. There are many published tools for the *de novo* identification of type III and type IV secreted proteins^3^. However, in our testing, none of these tools were consistent in identifying known effectors or were only available as webservers, and several tools identified many false positives that are unlikely to encode for secreted proteins. Therefore, we did not include the identification of T3SS and T4SS secreted proteins in the current iteration of Beav. Tools that can correctly identify secreted proteins with few false positives are needed for accurate annotation.

Other features to investigate for future Beav versions include annotation of other MGEs, such as transposons and plasmids, especially in draft genome assemblies. Comprehensive annotation of regulatory elements, such as promoters and transcription factor binding sites, is also a future development goal. Several programs exist for the prediction of bacterial promoters, such as the sigma70 promoter^82–84^. However, most promoter prediction tools were developed based on specific bacterial taxa and do not work well for analyses outside of closely related lineages. There is a need for *de novo* promoter annotation that captures the full breadth of promoters and regulatory elements across diverse bacteria.

## Acknowledgements

We would like to thank Danielle Stevens for testing the Beav pipeline and providing helpful feedback. We would also like to thank Oliver Schwengers for assisting in maintaining dependency compatibility with Bakta, and Björn Grüning for helpful advice on Bioconda integration. This work was funded by startup funds from the Department of Botany and Plant Pathology at Oregon State University.

